# Hypoxia reduces cell attachment of SARS-CoV-2 spike protein by modulating the expression of ACE2, neuropilin-1, syndecan-1 and cellular heparan sulfate

**DOI:** 10.1101/2021.01.09.426021

**Authors:** Endika Prieto-Fernández, Leire Egia-Mendikute, Laura Vila-Vecilla, Alexandre Bosch, Adrián Barreira-Manrique, So Young Lee, Ana García-del Río, Asier Antoñana-Vildosola, Borja Jiménez-Lasheras, Leire Moreno-Cugnon, Jesús Jiménez-Barbero, Edurne Berra, June Ereño-Orbea, Asis Palazon

**Affiliations:** Cancer Immunology and Immunotherapy Lab, Centre for Cooperative Research in Biosciences (CIC bioGUNE), Basque Research and Technology Alliance (BRTA), Bizkaia Technology Park, Building 801A, Derio, Spain; Cancer Cell Signaling and Metabolism Lab, Centre for Cooperative Research in Biosciences (CIC bioGUNE), Basque Research and Technology Alliance (BRTA), Bizkaia Technology Park, Building 801A, Derio, Spain; Chemical Glycobiology Lab, Centre for Cooperative Research in Biosciences (CIC bioGUNE), Basque Research and Technology Alliance (BRTA), Bizkaia Technology Park, Building 801A, Derio, Spain; Department of Organic Chemistry II, Faculty of Science and Technology, University of the Basque Country (UPV/EHU), Leioa, Spain; Ikerbasque, Basque Foundation for Science, Bilbao, Spain; CIBERONC (Centro de Investigación Biomédica en Red de Cáncer), Madrid, Spain

**Keywords:** SARS-CoV-2, Hypoxia, ACE2, Heparan sulfate, Syndecan-1

## Abstract

A main clinical parameter of COVID-19 pathophysiology is hypoxia. Here we show that hypoxia decreases the attachment of the receptor binding domain (RBD) and the S1 subunit (S1) of the spike protein of SARS-CoV-2 to epithelial cells. In Vero E6 cells, hypoxia reduces the protein levels of ACE2 and neuropilin-1 (NRP1), which might in part explain the observed reduction of the infection rate. In addition, hypoxia inhibits the binding of the spike to NCI-H460 human lung epithelial cells by decreasing the cell surface levels of heparan sulfate (HS), a known attachment receptor of SARS-CoV-2. This interaction is also reduced by lactoferrin, a glycoprotein that blocks HS moieties on the cell surface. The expression of syndecan-1, an HS-containing proteoglycan expressed in lung, is inhibited by hypoxia on a HIF-1α-dependent manner. Hypoxia or deletion of syndecan-1 results in reduced binding of the RBD to host cells. Our study indicates that hypoxia acts to prevent SARS-CoV-2 infection, suggesting that the hypoxia signaling pathway might offer therapeutic opportunities for the treatment of COVID-19.

## Introduction

By mid-May of 2021, SARS-CoV-2 has infected over 160 million people and caused >3.3 million deaths. Until now, more than 1200 million vaccines have been administered. However, the efficacy of available vaccines against emerging variants remains unclear [1, 2]. Therefore, the need of effective treatments to prevent the infection or to manage the disease in severe cases of COVID-19 remains an urgent medical need.

The spike protein of SARS-CoV-2 interacts with the angiotensin-converting enzyme 2 (ACE2), a key-receptor widely expressed on the respiratory tract, through the receptor binding domain (RBD) [3, 4, 5]. Other factors such as neuropilin-1 (NRP1) [6, 7] and the transmembrane protease serine 2 (TMPRSS2) [8] contribute to the entry of SARS-CoV-2 into the host cells. Another layer of interaction between the virus and the host that facilitates viral entry is the attachment of the spike protein to cellular heparan sulfate (HS) [5]. HS is part of the cellular glycocalyx and decorates the HS-containing proteoglycans (HSPGs). The family of syndecans are HSPGs expressed on several cell types and organs [9], being syndecan-1 the most relevant in terms of expression in the lung [10]. The interaction between the spike and HS can be efficiently blocked by lactoferrin [11].

A common feature of COVID-19 patients is moderate to severe hypoxia [12, 13, 14, 15]. Disease severity correlates with the level of naturally occurring anti-SARS-CoV-2 antibodies in serum [16], and patients with severe disease are characterized by low oxygen saturation that often results in the need of oxygen supplementation [17]. Currently, the impact of hypoxia on SARS-CoV-2 infectivity and COVID-19 pathogenesis is unclear, but there are emerging indications that low oxygen concentrations might prevent infection or disease severity [18, 19]. This notion is supported, in part, by the fact that epidemiological data indicates that COVID-19 severity decreases in high altitude areas [20].

There is evidence that hypoxia regulates the expression of ACE2, but the underlying mechanisms and impact on physiology and disease remain controversial. Previous studies have demonstrated that regulation of ACE2 expression by hypoxia is context specific and cell type- and time-dependent [21, 22, 23, 24, 25]. Elucidating these mechanisms in lung epithelial cells would help to understand the cellular susceptibility to infection by SARS-CoV-2 under limited oxygen availability. Additionally, the members of the syndecan family, the main cellular HSPGs, are also differentially regulated by hypoxia. For example, the expression of syndecan-1 in the lungs is reduced under hypoxia [26, 27].

Cell responses to hypoxia are mediated by the family of hypoxia inducible factors (HIF). COVID-19 disease leads to the accumulation of HIF in the lung, providing a molecular link between SARS-CoV-2 infection and hypoxia in patients with severe disease [28]. Our hypothesis is that hypoxia modulates the expression of entry receptors and attachment factors on host cells. In this context, elucidating the underlying mechanisms of this regulation could lead to the development of novel therapeutic strategies for the management of COVID-19, including those targeting the hypoxia response pathway.

## Materials and methods

### Cell Culture

The NCI-H460 human lung epithelial cell line was obtained from DSMZ (German Collection of Microorganisms and Cell Lines Cat#: ACC737) and cultured in RPMI medium 1640 GlutaMAX (Gibco Cat#: 61870044) according to standard culture protocols. Vero E6 monkey kidney cells (ATCC Cat#: CRL-1586) were kindly provided by Nicola G.A. Abrescia (CIC bioGUNE) and cultured in MEM (Gibco Cat#: 31095-029). HEK 293T cells (Takara Bio Inc. Cat#: 632180) were cultured in DMEM (Gibco Cat#: 41966-029). Media were supplemented with 10% FBS (Thermo Fisher Cat#: 10270106) and 1% Penicillin-Streptomycin (Thermo Fisher Cat#: 15140122). Hypoxia cultures were performed at 1% O_2_ and 5% CO_2_ in an In Vivo2 400 hypoxia station (Ruskinn Technologies).

### Lactoferrin assay

Binding of RBD to the surface of cells was measured by flow cytometry after incubation with increasing doses of human lactoferrin (0, 1, 5 and 10 mM) (Sigma Aldrich Cat#: L1294) in a final volume of 100 μL of culture medium. The incubation was done for 45 minutes in a cell incubator with standard conditions (37°C, 21% oxygen and 5% CO_2_).

### Assessment of binding of RBD and S1 to the cell surface

After culture, cells were collected with cell dissociation buffer (Thermo Fisher Cat#: 13151-014), distributed in 96-well polystyrene conical bottom plates (Thermo Fisher Cat#: 249570) and washed with PBS (Fisher BioReagents Cat#: BP3994) containing 0.5% BSA (Sigma Aldrich Cat#: A9647) (blocking buffer) followed by centrifugation and the supernatant was discarded. Cells were incubated with biotinylated S1 (Acrobiosystems Cat#: S1N-C82E8) or RBD (Acrobiosystems Cat#: SPD-C82E9) proteins (20 μg/mL) in 50 μL of blocking buffer for 30 minutes at 4°C. After incubation, cells were centrifuged, washed with 200 μL of blocking buffer, centrifuged again and incubated with streptavidin-PE (Thermo Fisher Cat#: 12-4317-87) diluted in 100 μL of blocking buffer (1:200 dilution). Finally, cells were centrifuged and washed twice and resuspended in blocking buffer with DAPI (Invitrogen Cat#: D1306) for acquisition in the flow cytometer. All centrifugation steps were performed at 300 x *g* for 5 min (4°C).

### Generation of pseudotyped lentiviral particles

Pseudotyped lentiviral particles were generated by transfecting HEK 293T cells as described in Crawford et al. [1], using a five-plasmid third generation system kindly provided by Dr. Jesse D. Bloom (Fred Hutchinson Cancer Research Center) and Dr. Jean-Philippe Julien (University of Toronto). Confluent HEK 293T cells (50-70%) were transfected with plasmids encoding for the lentiviral backbone (containing a CMV promoter to express ZsGreen) (NR-52520, 5.79 µg), the SARS-CoV-2 spike protein (NR-52514, 1.97 µg), HDM-Hgpm2 (NR-52517, 1.27 µg), pRC-CMV-Rev1b (NR-52519, 1.27 µg) and HDM-tat1b (NR-52518, 1.27 µg) using the jetPEI kit (Polyplus-transfection Cat#: 101-10N). 48 hours after transfection, viruses were harvested from the supernatant, filtered through a 0.45 µm filter (VWR Cat#: 514-0063), concentrated using Lenti-X Concentrator (Takara Bio Inc. Cat#: 631231) and stored in PBS at -80°C until use.

### Titration of the pseudotyped lentiviral particles and infection of Vero E6 cells

Viral titration was performed in Vero E6 cells as described in Crawford et al. [1]. For single-infection experiments, 150.000 cells were seeded in 6-well plates and cultured overnight in normoxia (21% O_2_) and incubated for additional 24 hours under normoxia or hypoxia (1% O_2_). After that, cells were incubated with the pseudotyped lentiviral particles (MOI: 0.1) for 48 hours under normoxic conditions. The infected cells expressed ZsGreen, allowing their detection by flow cytometry.

### Deletion of HIF-1α and syndecan-1 in NCI-H460 cells via CRISPR/Cas9

Deletion of HIF-1α in NCI-H460 cells was performed using a pool of two sgRNAs from Synthego (sgRNA sequences: gaugguaagccucaucacag and guuuuccaaacuccgacauu) and the TrueCut Cas9 protein v2 (Invitrogen Cat#: A36497). Deletion of syndecan-1 was performed using the following TrueGuide Synthetic sgRNAs from Invitrogen: CRISPR913712_SGM and CRISPR913714_SGM. RNP-complexes were generated with 7.5 pmol of Cas9, 7.5 pmol of the sgRNA-1 and 7.5 pmol of the sgRNA-2. RNP-complexes were introduced into cells using the Lipofectamine CRISPRMAX Cas9 Transfection reagent (Invitrogen Cat#: CMAX00001) and Opti-MEM reduced serum medium (Gibco Cat#: 31985062) following the Pub. No. MAN0017058 by Invitrogen (pages 5 and 6). A non-targeting sgRNA that does not recognize any sequence in the human genome was used as a negative control (Invitrogen Cat#: A35526). The deletion efficiency of HIF-1α and syndecan-1 proteins was measured by western blot and flow cytometry, respectively, and the pool of edited cells was used for downstream experiments.

### Flow cytometry

Cells were collected, washed with blocking buffer and incubated with fluorochrome-conjugated antibodies (1:100 dilution in blocking buffer) against syndecan-1 (BD Biosciences Cat#: 565943), syndecan-2 (R&D Systems Cat#: FAB2965P), syndecan-3 (R&D Systems Cat#: FAB3539A) and syndecan-4 (R&D Systems Cat#: FAB291819) for 15 minutes at 4°C. Total cellular heparan sulfate (HS) was measured by flow cytometry after incubation for 30 minutes at 4°C with an anti-HS primary antibody (Amsbio Cat#: 370255-S; clone F58-10E4) (1:200 dilution in blocking buffer). After that, cells were centrifuged, washed and incubated with an anti-IgM FITC-conjugated secondary antibody (Miltenyi Biotec Cat#: 130-095-906) (1:500 dilution in blocking buffer). Finally, cells were centrifuged, washed twice and resuspended in 200 μL of blocking buffer in the presence of 7-AAD (BD Biosciences Cat#: 51-68981E) or DAPI (Invitrogen Cat#: D1306) to discriminate alive cells. Cells were acquired on a FACSymphony cytometer (BD Biosciences). The results were analyzed using FlowJo version 10 (BD Biosciences).

### Quantitative PCR

Total RNA was isolated using the NucleoSpin RNA kit (Macherey-Nagel Cat#: 740955.250). The cDNA was synthesized by RT-PCR from 1 μg of purified RNA with the M-MLV reverse transcriptase (Thermo Fisher Cat#: 28025-013) and random primers (Thermo Fisher Cat#: 58875). The quantitative PCR (Q-PCR) reactions were conducted in triplicate on a ViiA 7 Real-Time PCR system (Thermo Fisher) from 1 μL of cDNA using the PerfeCTa SYBR Green SuperMix reagent (Quantabio Cat#: 95056-500) and gene-specific primers. The amplification program consisted of initial denaturation at 95°C for 3 min followed by 40 cycles of denaturation at 95°C for 15 s, annealing at 60°C for 60 s and extension at 72°C for 60 s. Data were analyzed using the QuantStudio software version 1.3 (Thermo Fisher). The relative quantification in gene expression was determined using the 2^−ΔΔCt^ method by using the *RPLP0* gene as a housekeeping gene (forward: 5’-CGACCTGGAAGTCCAACTAC-3’ and reverse: 5’-ATCTGCTGCATCTGCTTG-3’). *PGK1*, a HIF-1α target gene, was used as a control for hypoxia (forward: 5’-CCGCTTTCATGTGGAGGAAGAAG-3’ and reverse: 5’-CTCTGTGAGCAGTGCCAAAAGC-3’). Primer sequences for measuring *ACE2* and *SDC1* expression on human NCI-H460 cells were previously described by Ma et al. [2] and Reynolds et al. [3], respectively. As these oligos presented several mismatches with the corresponding genes on the reference genome of the African green monkey, we designed the following primers to amplify *ACE2* from Vero E6 cells (forward: 5’-AGCACTCACGATTGTTGGGA-3’ and reverse: 5’-CCACCCCAACTATCTCTCGC-3’).

### Western Blot

Total cell lysates were collected using RIPA buffer and quantified with the BCA Protein Assay kit (Thermo Fisher Cat#: 23227). Samples were mixed with LDS sample buffer (Invitrogen Cat#: NP0007) containing DTT, boiled for 15 minutes, separated in a 4-15% Mini-PROTEAN TGX precast protein gels (BioRad Cat#: 4561083) and transferred to a 0.2 μm PVDF membranes (BioRad Cat#: 1704156) using a Trans-Blot Turbo transfer system (BioRad). PageRuler protein ladder (Thermo Fisher Cat#: 26619) was used to estimate the molecular weight of the proteins. The membranes were blocked for 1 hour in 5% skim milk (Millipore Cat#: 70166) and 0.5% Tween-20 (Sigma Aldrich Cat#: 9005-64-5) diluted in PBS. The membranes were probed overnight at 4°C with primary antibodies diluted in PBS containing 5% BSA and 0.5% Tween-20, washed five times with PBS (containing 0.5% Tween-20), incubated for 1 hour at RT with the corresponding secondary HRP-conjugated antibodies (1:5000 diluted in 5% skim milk and 0.5% Tween-20 diluted in PBS) and washed for five additional times. The membranes were probed with antibodies against HIF-1α (Novus Biologicals Cat#: NB100-449), ACE2 (Bioss antibodies Cat#: bsm-52614R), NRP1 (Novus Biologicals Cat#: NBP2-67539), TMPRSS2 (Novus Biologicals Cat#: NBP3-00492), β-actin (Cell Signaling Cat#: D6A8) and β-tubulin (Thermo Fisher Cat#: MA5-16308). The HRP-conjugated secondary antibodies against mouse (Cat#: S301677076S) and rabbit (Cat#: S301677074S) were obtained from Cell Signaling. Chemiluminescence detection was performed using Clarity Max Western ECL Substrate (BioRad Cat#: 170506) on an iBright system (Invitrogen). Band densitometry was performed using ImageJ (https://imagej.nih.gov/ij/).

### Statistics

Statistical analyses were performed using GraphPad Prism version 8.0. The test applied in each panel is specified on the figure legends.

## Results

### Hypoxia reduces the binding of the SARS-CoV-2 spike to epithelial cells

In order to understand if hypoxia could affect the cellular binding capacity of the SARS-CoV-2 spike protein, we subjected Vero E6 cells or human lung epithelial cells (NCI-H460) to different oxygen concentrations (21% O_2_ or 1% O_2_) and measured the binding ability of the receptor binding domain (RBD) and the S1 subunit (S1) of the spike protein to the surface of these cells in vitro. Figure 1 shows a significant decrease of the binding of RBD and S1 after culturing Vero E6 (Figure 1a) or NCI-H460 (Figure 1b) cells under hypoxia (1% O_2_) compared to normoxia (21% O_2_).

**Figure 1.**
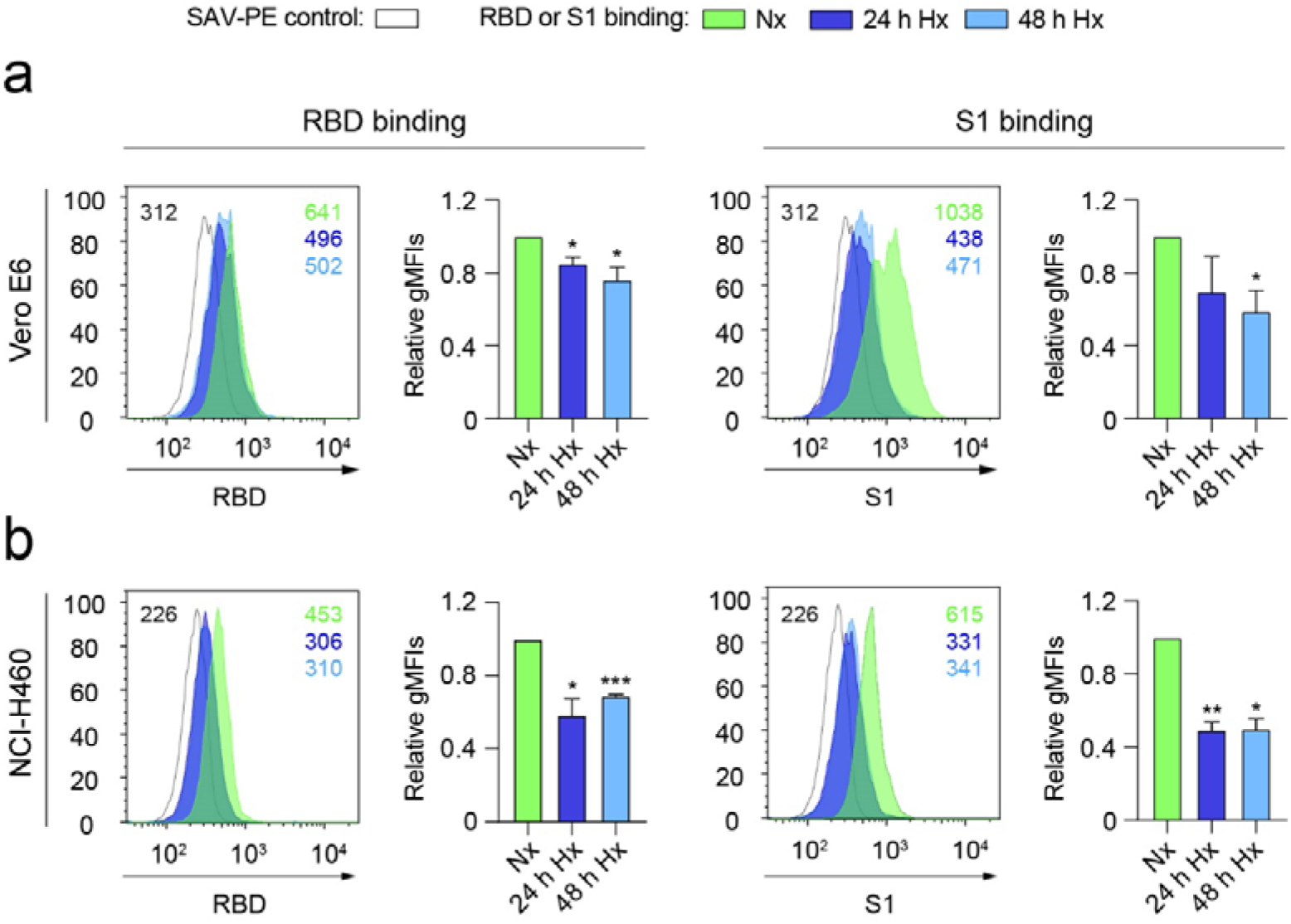
Hypoxia reduces the binding of the SARS-CoV-2 spike to epithelial cells. (*a, b*) Binding of the receptor binding domain (RBD) (*left*) or S1 subunit (S1) (right) to Vero E6 (*a*) (n=3 independent experiments, unpaired t test) or NCI-H460 (*b*) (n=2 independent experiments, unpaired t test) cells cultured under normoxia (21% O_2_) or hypoxia (1% O_2_) for 24 and 48 hours, measured by flow cytometry. A representative histogram indicating the geometric mean fluorescent intensity (gMFI) value for each condition is shown. Bar graphs represent RBD or S1 binding relative to normoxia. Error bars represent SEM and asterisks represent p values (*, ≤ 0.05; **, < 0.01; ***, < 0.001). Abbreviations: SAV-PE: Streptavidin-phycoerythrin; Nx: normoxia (21% O_2_); Hx: hypoxia (1% O_2_).

### Hypoxia decreases ACE2 and NRP1 protein levels on Vero E6 cells

To further explore the mechanism underlying the observed reduced binding of the spike under hypoxia, we assessed ACE2 gene and protein expression on Vero E6 and NCI-H460 cells by Q-PCR and western blot, respectively. Vero E6 cells, but not NCI-H460 cells, present detectable levels of ACE2 transcripts (Figure 2a). Accordingly, ACE2 protein is only detectable on Vero E6 cells (Figure 2b), suggesting that the observed binding of the spike to the surface of NCI-H460 cells is ACE2-independent. We then explored if hypoxia modulates ACE2 expression on Vero E6 cells. Hypoxia culture results in a transient accumulation of HIF-1α and upregulation of the expression of ACE2 (Figure 2b and 2c). After 48 hours of exposure to limited oxygen supply, the level of ACE2 protein on Vero E6 cells is significantly reduced (Figure 2b). On a similar manner, we measured the expression of other important factor that mediate the initial steps of SARS-CoV-2 infection: NRP1 and TMPRSS2. NRP1 (but not TMPRSS2) is significantly reduced on Vero E6 cells after culture for 48 hours under hypoxia (Figure 2d), as previously described by Casazza et al. [32] on the hypoxic tumor microenvironment. NCI-H460 cells do not express detectable levels of NRP1.

**Figure 2.**
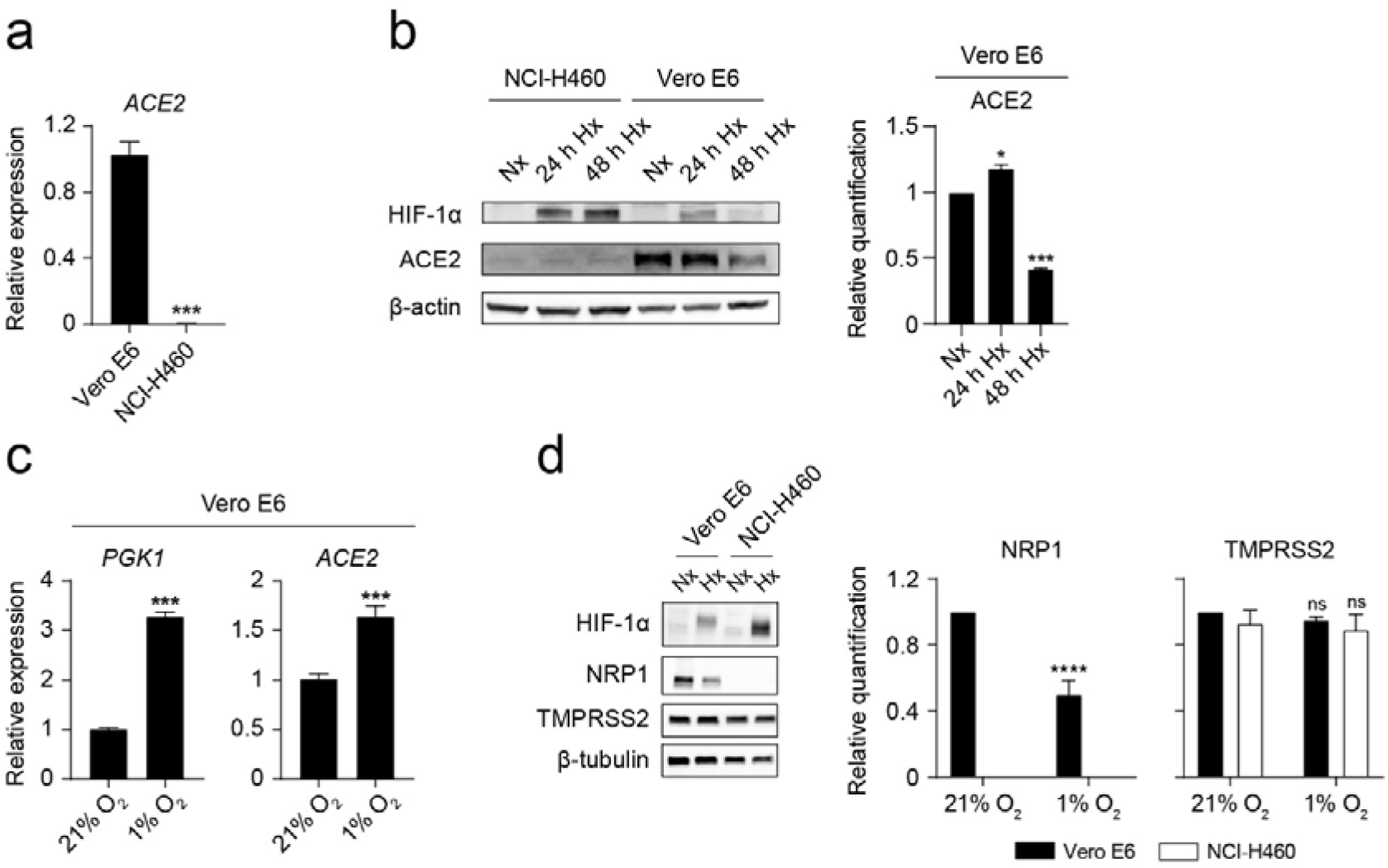
Hypoxia decreases ACE2 and NRP1 protein levels on Vero E6 cells. (*a*) Relative *ACE2* gene expression on Vero E6 and NCI-H460 measured by Q-PCR (n=3, unpaired t test). (*b*) (Left) Western blot of HIF-1α, ACE2 and β-actin on NCI-H460 and Vero E6 cells cultured under normoxia (21% oxygen) or hypoxia (1% oxygen) for the indicated time points. (Right) Relative quantification of ACE2 protein expression by densitometry (n=2, one-way ANOVA). (*c*) Relative gene expression of *PGK1* (left) and *ACE2* (right) on Vero E6 cells cultured under normoxia or hypoxia for 24 hours (n=3, unpaired t test). (*d*) (Left) Western blot of HIF-1α, NRP1, TMPRSS2 and β-tubulin on Vero E6 and NCI-H460 cells cultured under normoxia or hypoxia for 48 hours. (Right) Relative quantification of NRP1 and TMPRSS2 proteins by densitometry (n=3, 2-way ANOVA). Error bars represent SEM and asterisks represent p values (*, ≤ 0.05; ***, < 0.001; ****, <0.0001).

### Hypoxia prevents the infection of Vero E6 cells with pseudotyped viral particles expressing the spike protein of SARS-CoV-2

We then assessed the influence of oxygen availability on the infection capacity of pseudotyped lentiviral particles that express the full-length spike of SARS-CoV-2. Vero E6 cells exposed to hypoxia show a significant decrease on the rate of infection (Figure 3a) compared to normoxia. NCI-H460 cells, which lack expression of ACE2 and NRP1 (Figures 2b and 2d), are not amenable to infection (Figure 3b).

**Figure 3.**
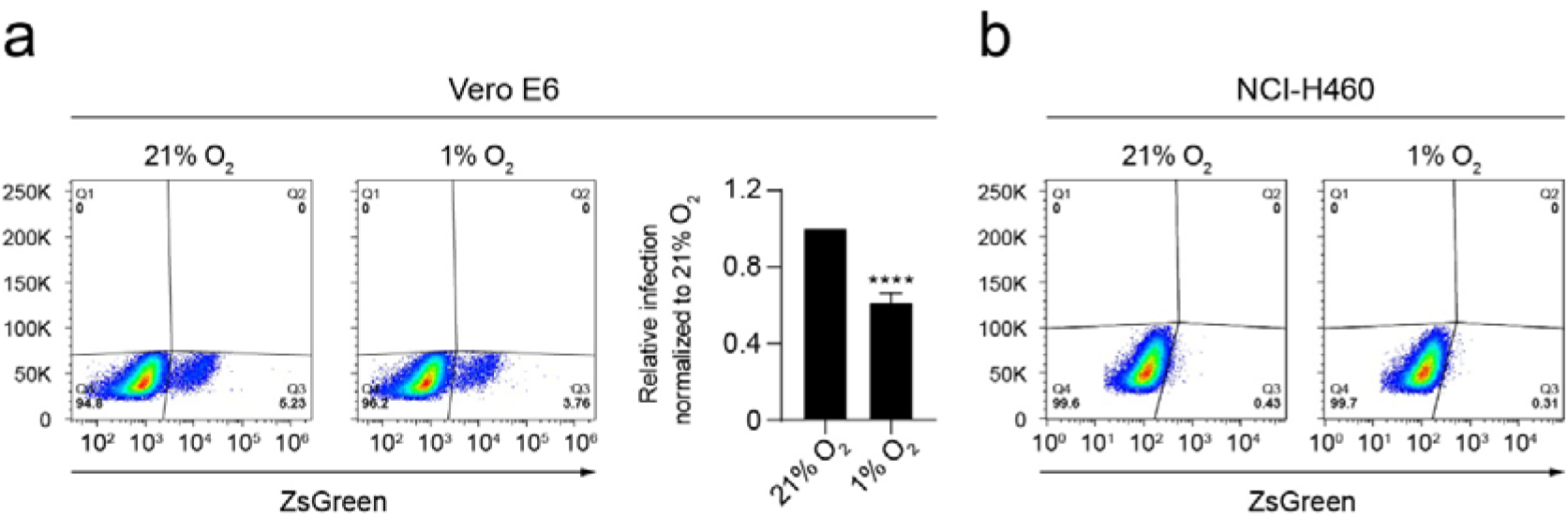
Hypoxia prevents the infection of Vero E6 cells with pseudotyped viral particles expressing the spike protein of SARS-CoV-2. (*a*) Dot plots representing the infection rate of Vero E6 cells exposed to normoxia (21% O_2_) or hypoxia (1% O_2_) measured by flow cytometry (ZsGreen expression) 48 hours after addition of the pseudotyped viral particles (n=6 independent experiments, unpaired t test). Gating strategy was based on uninfected cells. Bar graphs represent infectivity relative to normoxia. (*b*) NCI-H460 cells are not infected by pseudotyped viral particles. Error bars represent SEM and asterisks represent p values (****, p< 0.0001).

### Lactoferrin blocks the binding of RBD to the surface of epithelial cells

Recently, it has been demonstrated that the spike of SARS-CoV-2 can bind to HS expressed on the host cell surfaces [5]. We reasoned that this phenomenon might explain the observed binding of RBD to NCI-H460 epithelial cells in the absence of ACE2 and NRP1. To confirm this interaction, we blocked cellular HS with human lactoferrin, a natural glycoprotein that binds to HS and presents antiviral activity against coronaviruses [11, 33]. Figure 4 shows that lactoferrin was able to reduce the attachment of the RBD of SARS-CoV-2 to Vero E6 (Figure 4a) and NCI-H460 (Figure 4b) cells on a dose-dependent manner. Interestingly, the degree of blockade by lower doses of lactoferrin was higher on NCI-H460 compared to Vero E6 cells.

**Figure 4.**
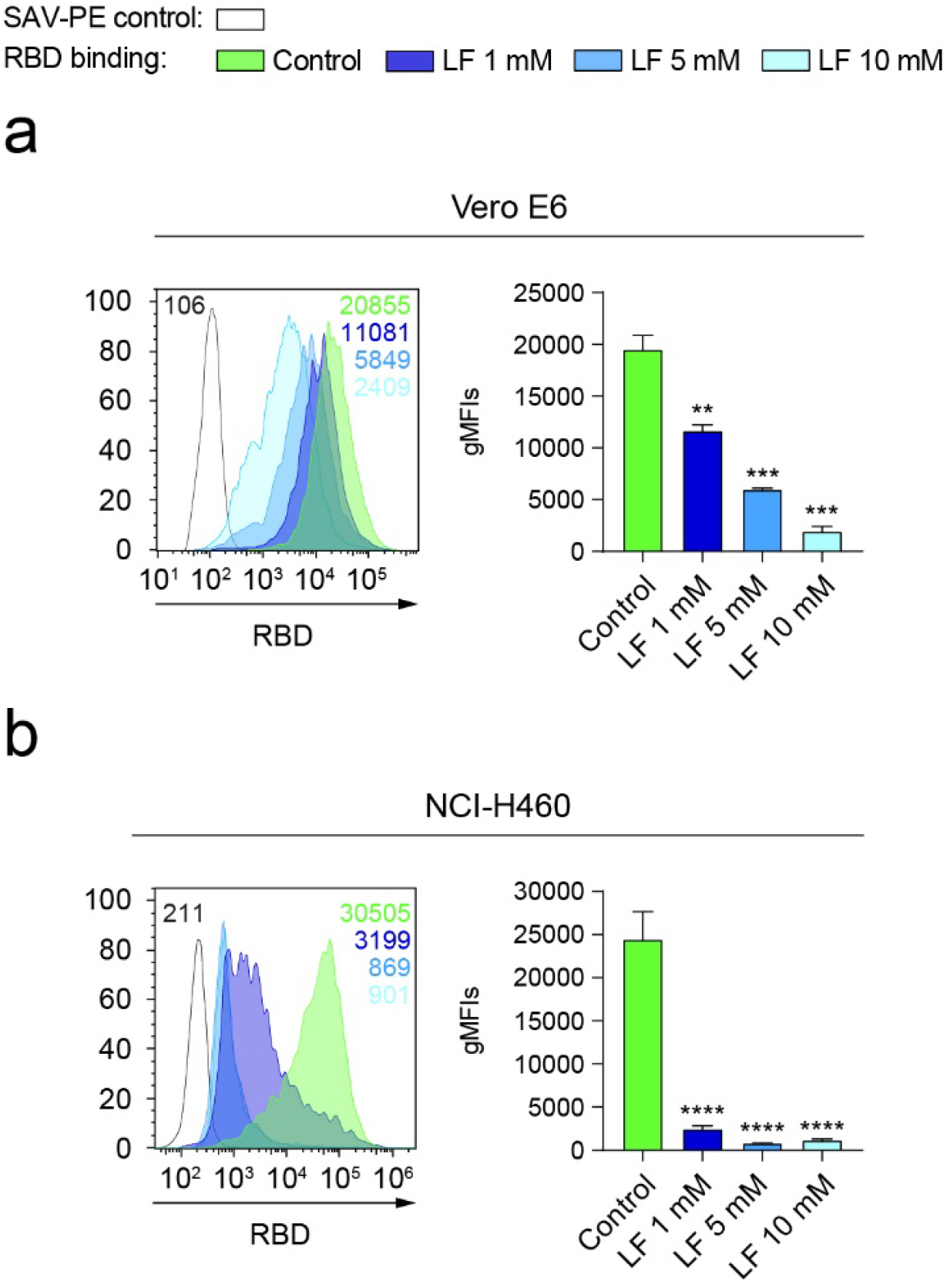
Lactoferrin blocks the binding of RBD to the surface of epithelial cells. (*a, b*) Binding of RBD to Vero E6 (*a*) (n=2 independent experiments, one-way ANOVA) or NCI-H460 (*b*) (n=3 independent experiments, one-way ANOVA) cells cultured under normoxia (21% O_2_) for 48 hours measured by flow cytometry after incubation with increasing doses of lactoferrin (LF). A representative flow cytometry histogram indicating the gMFI value for each condition is shown. Error bars represent SEM and asterisks represent p values (**, < 0.01; ***, < 0.001; ****, < 0.0001).

### Hypoxia decreases the total level of heparan sulfate on the surface of epithelial cells

Considering that cellular HS mediates the binding of RBD in NCI-H460 cells lacking the expression of ACE2 and NRP1, we explored if this interaction was altered by hypoxia. Hypoxia significantly decreases the total level of cellular HS present on Vero E6 cells (Figure 5a) and, to a higher extent, on NCI-H460 cells (Figure 5b). The syndecan family is a major contributor to the pool of HS on the surface of cells. In this context, we assessed the expression levels of syndecan-1, syndecan-2, syndecan-3 and syndecan-4 on NCI-H460 cells. We found that syndecan-1 and syndecan-3 are the main syndecans expressed on this cell model and that hypoxia significantly reduces their expression at protein level (Figure 5c).

**Figure 5.**
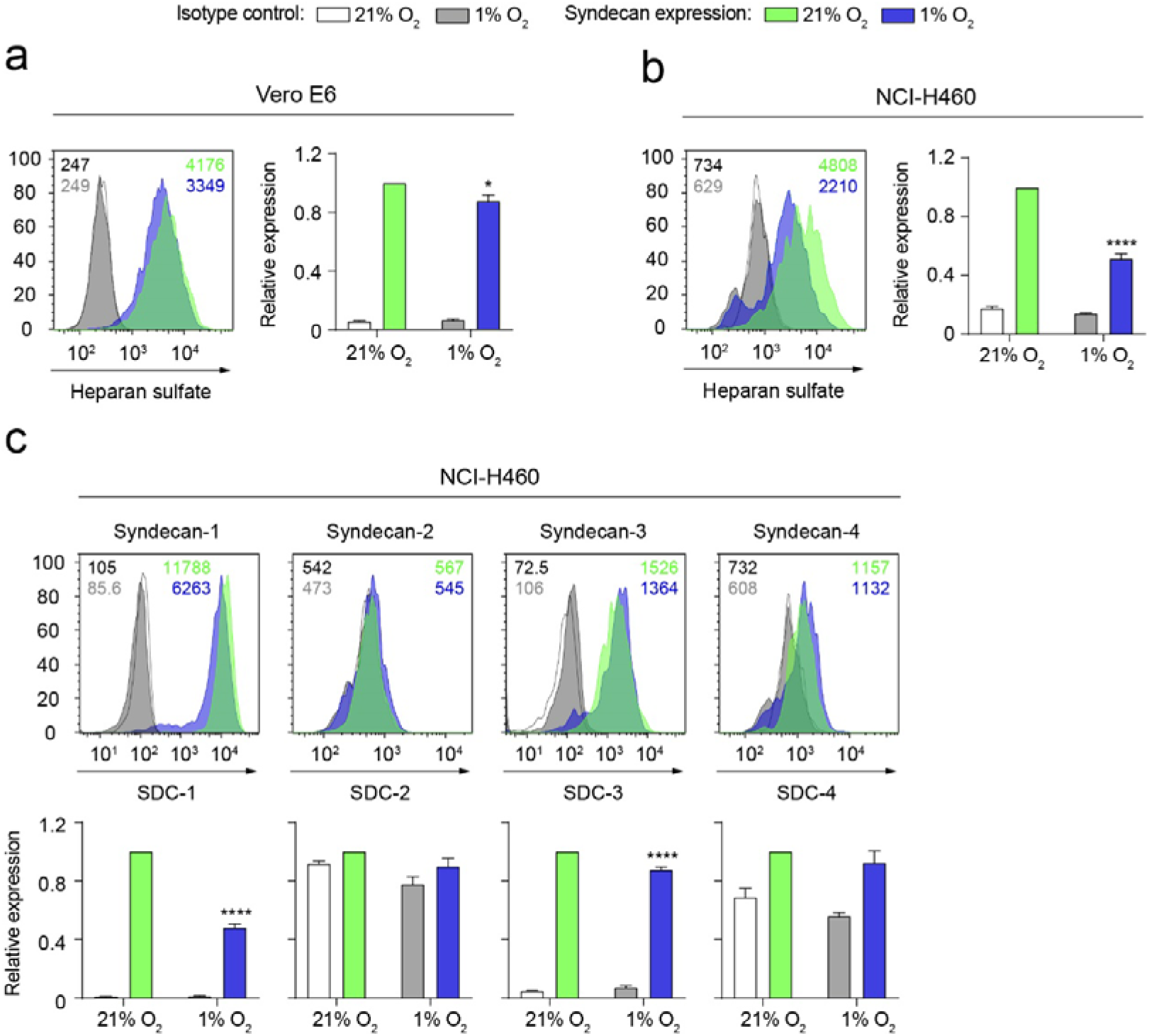
Hypoxia decreases the total level of heparan sulfate on the surface of epithelial cells. (*a, b*) Total heparan sulfate (HS) levels on Vero E6 (*a*) and NCI-H460 (*b*) cells cultured under normoxia (21% O_2_) or hypoxia (1% O_2_) for 24 hours, measured by flow cytometry. (*c*) Expression of the syndecan family members on NCI-H460 cells cultured under normoxia or hypoxia for 24 hours. Representative flow cytometry histograms including gMFI values are shown. Bar graphs represent syndecan expression levels under hypoxia relative to normoxia (n=2 independent experiments, 2-way ANOVA). Error bars represent SEM and asterisks represent p values (*, ≤ 0.05; ****, ≤ 0.0001).

### Hypoxia reduces the binding of RBD to NCI-H460 epithelial cells by downregulating syndecan-1 expression on a HIF-1α-dependent mechanism

Based on the finding that syndecan-1 is the most abundantly expressed syndecan on NCI-H460 cells (Figure 5c), we sought to ascertain its role on the attachment of the spike protein to host cells. Figure 6a shows that NCI-H460 cells expressing high levels of syndecan-1 capture higher amounts of RBD compared to cells expressing low levels of syndecan-1. To further demonstrate the binding of RBD to syndecan-1, we generated a cell line with reduced expression of syndecan-1 via CRISPR/Cas9 (SDC1-KO). A pool of SDC1-edited cells binds less RBD compared to control NCI-H460 cells (Figures 6b). Given that HIF-1α is the main transcriptional mediator of the response to hypoxia, we checked if it was responsible for the observed downregulation of syndecan-1 under hypoxia (Figure 5c and 6c). To this end, we generated HIF-1α deficient NCI-H460 cells via CRISPR/Cas9 (Figure 6d). Deletion of HIF-1α on NCI-H460 cells partially rescued the observed inhibition of syndecan-1 expression under hypoxia (Figure 6e), resulting in a marked increase in the total levels of cellular HS (Figure 6f). Consequently, the level of RBD binding to HIF1-KO cells under hypoxia was higher than in control HIF-1α competent cells (Figure 6g).

**Figure 6.**
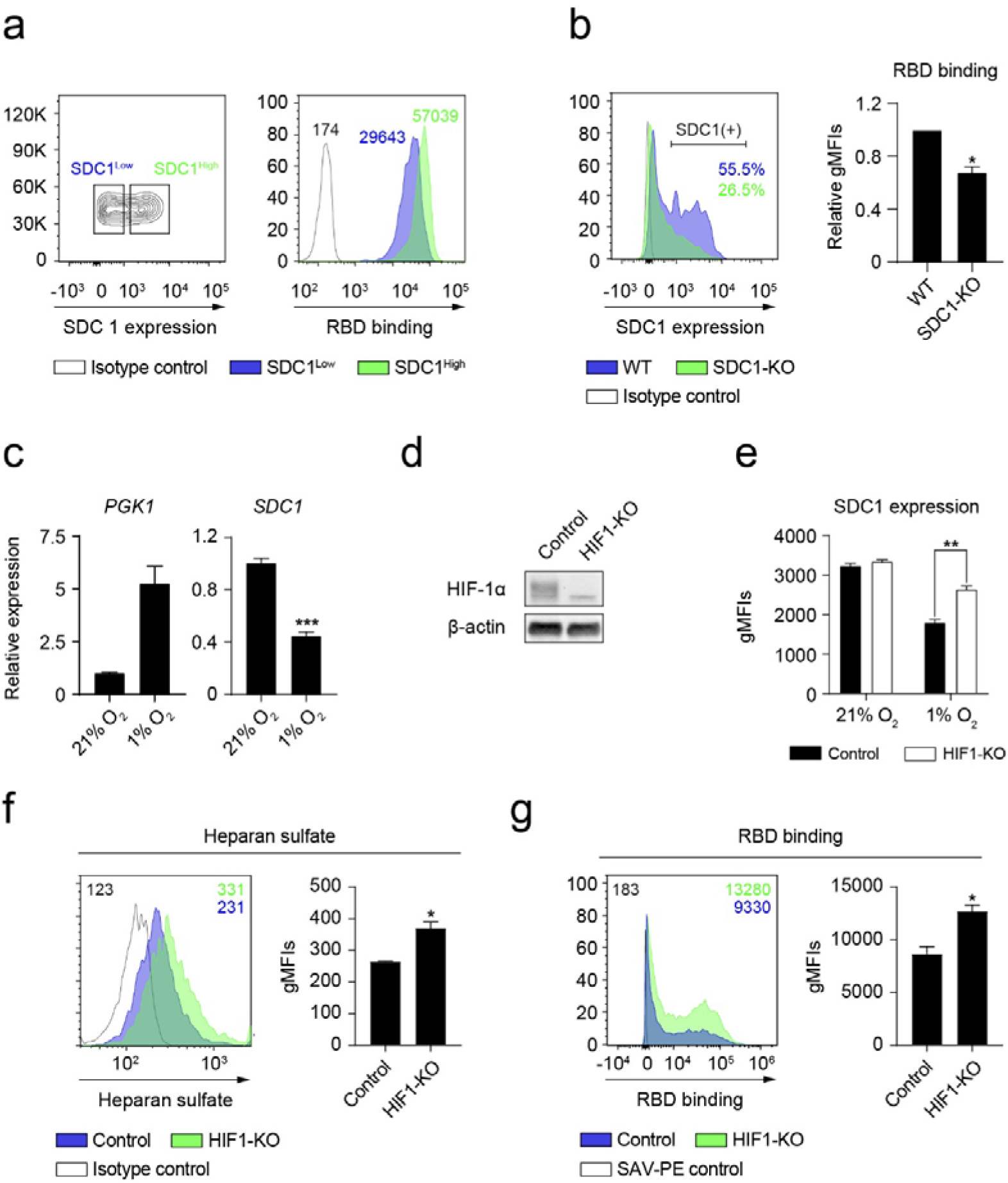
Hypoxia reduces the binding of RBD to NCI-H460 epithelial cells by downregulating syndecan-1 expression on a HIF-1α dependent mechanism. (*a*) (Left) Gating strategy followed to select cells expressing high (SDC1^High^) or low (SDC1^Low^) levels of syndecan-1 under normoxia. (Right) Representative flow cytometry histogram showing the binding of RBD to SDC1^High^ (green) or SDC1^Low^ (blue) cell populations (n=2 independent experiments). gMFI values are shown. (*b*) (Left) A representative histogram showing the percentage of cells expressing syndecan-1 in the SDC1-KO or control cells. (Right) Binding of RBD to SDC1-KO cells normalized to control cells (n=2 independent experiments). (*c*) Relative gene expression of *PGK1* (left) and *SDC1* (right) on NCI-H460 cells cultured under normoxia or hypoxia for 24 hours (n=3, unpaired t test). (*d*) Western blot showing the lack of HIF-1α protein in HIF1-KO NCI-H460 cells cultured under hypoxia for 24 hours. (*e*) Syndecan-1 expression on HIF1-KO NCI-H460 cells cultured under normoxia or hypoxia for 24 hours, gMFI values are shown. (*f, g*) Total heparan sulfate level (*f*) and RBD binding (*g*) on HIF1-KO NCI-H460 cells (n=2 independent experiments, unpaired t test) cultured under hypoxia for 24 hours, gMFI values are shown. Representative flow cytometry histograms including gMFI values are shown. Error bars represent SEM and asterisks represent p values (*, ≤ 0.05; **, < 0.01; ***, < 0.001).

## Discussion

COVID-19 disease is associated with hypoxia [12, 28, 34], which results on decreased oxygen availability for tissues. This phenomenon affects the susceptibility of host cells to viral infections [35] and shapes immune responses [36]. Here we describe the binding ability of the spike of SARS-CoV-2 to epithelial cells cultured under different oxygen concentrations. We show that hypoxia inhibits the binding of the RBD and the S1 subunit of the spike of SARS-CoV-2 to host cells by modulating the expression of several entry and attachment factors.

In this study we consider two different and complementary cellular models. First, Vero E6 cells, a widely used cell type in SARS-CoV-2 biology research because of its expression of ACE2, NRP1 and TMPRSS2 factors, which are components of the cellular machinery that allow the entry of SARS-CoV-2 into host target cells. Second, NCI-H460 human lung epithelial cells, which lack the expression of ACE2 and NRP1 and as such, do not get infected by pseudotyped viral particles expressing the spike of SARS-CoV-2, but permits the binding of the spike and RBD to cellular HS [11, 37]. This dual approach allowed us to investigate the mechanism underlying the observed reduction of RBD binding and infectivity.

Using the Vero E6 model we showed that prolonged hypoxia reduces the expression levels of cell surface ACE2 and NRP1, resulting in a reduced susceptibility to infection by pseudotyped viral particles expressing the spike of SARS-CoV-2. We have not observed changes in the expression of TMPRSS2 under hypoxia. Previous studies suggest that the expression of ACE2 under hypoxia is context dependent [21, 22, 23, 24, 25]. Hypoxia exposure time is one of the variables that influence ACE2 levels. We show that acute hypoxia (24 hours) can induce a transient increase on ACE2 expression, probably mediated by the binding of HIF-1α to the hypoxia response elements found in the ACE2 promoter [22, 38]. However, chronic hypoxia (48 hours) leads to a significant reduction of ACE2 and HIF-1α levels on Vero E6 cells. In line with these findings, a recent study shows that hypoxia reduces the protein levels of ACE2 on several cell lines [38].

Cellular HS is known to cooperate with the entry receptor ACE2 as an attachment factor, facilitating the initial capture of viral particles by the cellular glycocalyx [5]. We used NCI-H460 human lung epithelial cells to disentangle the contribution of cellular HS to the observed reduction of infection under hypoxia on an ACE2-independent manner.

First, we confirmed that the RBD binds to HS expressed on target cells by blocking this interaction with lactoferrin. Vero E6 cells are less sensitive than NCI-H460 cells to the blockade exerted by low doses of lactoferrin, difference that could be attributed to the interaction of RBD with other cellular components that are absent in NCI-H460 cells, such as ACE2 and NRP1. These findings prompted us to use NCI-H460 cells as a model to assess the effect of hypoxia on the levels of cellular HS and its interaction with the spike of SARS-CoV-2.

We showed that hypoxia leads to a significant reduction of the cellular pool of HS on NCI-H460 cells. Interestingly, syndecan-1, which is expressed in the lung [10], is responsible for this decrease under hypoxia. We confirmed this finding by measuring the binding of RBD to cells expressing different amounts of syndecan-1. We then wanted to ascertain if HIF-1α was responsible for the reduction of the expression of syndecan-1 observed under hypoxia at both transcriptional and protein expression levels. Ablation of HIF-1α in NCI-H460 cells demonstrates that HIF-1α is necessary for the inhibition of syndecan-1 expression under hypoxia. Moreover, deletion of HIF-1α results in an increase of cellular HS levels and higher binding capacity of RBD to hypoxic NCI-H460 cells.

Together, these results indicate that the hypoxia response governed by HIF-1α in the host target cells leads to a protective effect against SARS-CoV-2 infection. In this context, the administration of HIF prolyl-hydroxylase (PHD) inhibitors that result in accumulation of HIF-1α may serve as a potential treatment to limit the initial attachment and entry of the virus into host cells [38, 39]. It should be noted that as the infection progresses into COVID-19 disease, exacerbated inflammatory responses contribute to severity. Inflammation and the response to hypoxia are closely related processes that coexists in certain microenvironments [40, 41]. Macrophages are important mediators of lung inflammation during COVID-19 disease [42]. As a result of hypoxia in the lung, elevated HIF-1α activity on immune innate cells contribute to severity [43, 44]. In this scenario, promoting systemic HIF-1α activity through pharmacological intervention in severe COVID-19 patients might be detrimental.

In summary, in this study we show that hypoxia decreases the binding of the spike of SARS-CoV-2 to epithelial cells by at least two different mechanisms: 1) decreasing the level of ACE2 and NRP1 expression; and 2) reducing the total amount of cellular HS (attachment factor) and syndecan-1 on the cell surface of epithelial cells.

## Limitations of the study

All experiments presented here have been performed in vitro with immortalized cell lines. The significance of these findings at physiopathological oxygen levels in vivo remains unexplored. In this context, the use of humanised mouse models and real SARS-CoV-2 virus would reinforce the findings of this study.

## Acknowledgements

This research was supported by the SPRI I+D COVID-19 fund (Basque Government, bG-COVID-19), the European Research Council (ERC) (grant numbers: ERC-2018-StG 804236-NEXTGEN-IO to A.P and ERC-2017-AdG 788143-RECGLYCANMR to J.J-B.), the Severo Ochoa Excellence Accreditation from MCIU (SEV-2016-0644) and the FERO Foundation. Personal fellowships: E.P. (Juan de la Cierva-Formación, FJC2018-035449-I), L.V. (Juan de la Cierva-Formación, FJCI-2017-34099), A.B. (AECC Bizkaia Scientific Foundation, PRDVZ19003BOSC), A.G. (Programa Bikaintek from the Basque Government, 48-AF-W1-2019-00012), A.A (La Caixa Inphinit, LCF/BQ/DR20/11790022), B.J. (Basque Government, PRE_2019_1_0320), L.M. (Juan de la Cierva-Formación, FJC2019-039983-I), E.B. (MINECO, (BFU2016-76872-R; Excellence Networks, SAF2017-90794-REDT) and A.P. (Ramón y Cajal, RYC2018-024183-I; Proyectos I+D+I, PID2019-107956RA-I00; and Ikerbasque Research Associate). The plasmids for the generation of pseudotyped lentiviral particles were kindly provided by Dr. Jesse D. Bloom (Fred Hutchinson Cancer Research Center) and Dr. Jean-Philippe Julien (The Hospital for Sick Children).

## References

1. Abdool Karim SS, de Oliveira T. New SARS-CoV-2 Variants - Clinical, Public Health, and Vaccine Implications. N Engl J Med. 2021 Mar 24. doi: 10.1056/NEJMc2100362. PubMed PMID: 33761203; PubMed Central PMCID: PMCPMC8008749.

2. Wang GL, Wang ZY, Duan LJ, et al. Susceptibility of Circulating SARS-CoV-2 Variants to Neutralization. N Engl J Med. 2021 Apr 6. doi: 10.1056/NEJMc2103022. PubMed PMID: 33822491.

3. Yan R, Zhang Y, Li Y, et al. Structural basis for the recognition of SARS-CoV-2 by full-length human ACE2. Science. 2020 Mar 27;367(6485):1444–1448. doi: 10.1126/science.abb2762. PubMed PMID: 32132184; PubMed Central PMCID: PMCPMC7164635.

4. Walls AC, Park YJ, Tortorici MA, et al. Structure, Function, and Antigenicity of the SARS-CoV-2 Spike Glycoprotein. Cell. 2020 Apr 16;181(2):281–292 e6. doi: 10.1016/j.cell.2020.02.058. PubMed PMID: 32155444; PubMed Central PMCID: PMCPMC7102599.

5. Clausen TM, Sandoval DR, Spliid CB, et al. SARS-CoV-2 Infection Depends on Cellular Heparan Sulfate and ACE2. Cell. 2020 Nov 12;183(4):1043–1057 e15. doi: 10.1016/j.cell.2020.09.033. PubMed PMID: 32970989; PubMed Central PMCID: PMCPMC7489987.

6. Cantuti-Castelvetri L, Ojha R, Pedro LD, et al. Neuropilin-1 facilitates SARS-CoV-2 cell entry and infectivity. Science. 2020 Nov 13;370(6518):856–860. doi: 10.1126/science.abd2985. PubMed PMID: 33082293; PubMed Central PMCID: PMCPMC7857391.

7. Daly JL, Simonetti B, Klein K, et al. Neuropilin-1 is a host factor for SARS-CoV-2 infection. Science. 2020 Nov 13;370(6518):861–865. doi: 10.1126/science.abd3072. PubMed PMID: 33082294.

8. Hoffmann M, Kleine-Weber H, Schroeder S, et al. SARS-CoV-2 Cell Entry Depends on ACE2 and TMPRSS2 and Is Blocked by a Clinically Proven Protease Inhibitor. Cell. 2020 Apr 16;181(2):271–280 e8. doi: 10.1016/j.cell.2020.02.052. PubMed PMID: 32142651; PubMed Central PMCID: PMCPMC7102627.

9. Sarrazin S, Lamanna WC, Esko JD. Heparan sulfate proteoglycans. Cold Spring Harb Perspect Biol. 2011 Jul 1;3(7). doi: 10.1101/cshperspect.a004952. PubMed PMID: 21690215; PubMed Central PMCID: PMCPMC3119907.

10. Parimon T, Yao C, Habiel DM, et al. Syndecan-1 promotes lung fibrosis by regulating epithelial reprogramming through extracellular vesicles. JCI Insight. 2019 Aug 8;5. doi: 10.1172/jci.insight.129359. PubMed PMID: 31393853; PubMed Central PMCID: PMCPMC6777916.

11. Hu Y, Meng X, Zhang F, et al. The in vitro antiviral activity of lactoferrin against common human coronaviruses and SARS-CoV-2 is mediated by targeting the heparan sulfate co-receptor. Emerg Microbes Infect. 2021 Dec;10(1):317–330. doi: 10.1080/22221751.2021.1888660. PubMed PMID: 33560940; PubMed Central PMCID: PMCPMC7919907.

12. Caputo ND, Strayer RJ, Levitan R. Early Self-Proning in Awake, Non-intubated Patients in the Emergency Department: A Single ED’s Experience During the COVID-19 Pandemic. Acad Emerg Med. 2020 May;27(5):375–378. doi: 10.1111/acem.13994. PubMed PMID: 32320506; PubMed Central PMCID: PMCPMC7264594.

13. Berenguer J, Ryan P, Rodriguez-Bano J, et al. Characteristics and predictors of death among 4035 consecutively hospitalized patients with COVID-19 in Spain. Clin Microbiol Infect. 2020 Nov;26(11):1525–1536. doi: 10.1016/j.cmi.2020.07.024. PubMed PMID: 32758659; PubMed Central PMCID: PMCPMC7399713.

14. Petrilli CM, Jones SA, Yang J, et al. Factors associated with hospital admission and critical illness among 5279 people with coronavirus disease 2019 in New York City: prospective cohort study. BMJ. 2020 May 22;369:m1966. doi: 10.1136/bmj.m1966. PubMed PMID: 32444366; PubMed Central PMCID: PMCPMC7243801 at www.icmje.org/coi_disclosure.pdf and declare: support from the Kenneth C Griffin Charitable Fund for submitted work; no financial relationships with any organizations that might have an interest in the submitted work in the previous three years; no other relationships or activities that could appear to have influenced the submitted work.

15. Yadaw AS, Li YC, Bose S, et al. Clinical features of COVID-19 mortality: development and validation of a clinical prediction model. Lancet Digit Health. 2020 Oct;2(10):e516–e525. doi: 10.1016/S2589-7500(20)30217-X. PubMed PMID: 32984797; PubMed Central PMCID: PMCPMC7508513.

16. Egia-Mendikute L, Bosch A, Prieto-Fernández E, et al. Sensitive detection of SARS-CoV-2 seroconversion by flow cytometry reveals the presence of nucleoprotein-reactive antibodies in unexposed individuals. Commun Biol. 2021 Apr 20;4(1):486. doi: 10.1038/s42003-021-02011-6. PubMed PMID: 33879833; eng.

17. Berlin DA, Gulick RM, Martinez FJ. Severe Covid-19. N Engl J Med. 2020 Dec 17;383(25):2451–2460. doi: 10.1056/NEJMcp2009575. PubMed PMID: 32412710.

18. Afsar B, Kanbay M, Afsar RE. Hypoxia inducible factor-1 protects against COVID-19: A hypothesis. Med Hypoin vitros. 2020 Oct;143:109857. doi: 10.1016/j.mehy.2020.109857. PubMed PMID: 32464493; PubMed Central PMCID: PMCPMC7238987 competing financial interests or personal relationships that could have appeared to influence the work reported in this paper.

19. Couzin-Frankel J. The mystery of the pandemic’s ‘happy hypoxia’. Science. 2020 May 1;368(6490):455–456. doi: 10.1126/science.368.6490.455. PubMed PMID: 32355007.

20. Arias-Reyes C, Zubieta-DeUrioste N, Poma-Machicao L, et al. Does the pathogenesis of SARS-CoV-2 virus decrease at high-altitude? Respir Physiol Neurobiol. 2020 Jun;277:103443. doi: 10.1016/j.resp.2020.103443. PubMed PMID: 32333993; PubMed Central PMCID: PMCPMC7175867.

21. Paizis G, Tikellis C, Cooper ME, et al. Chronic liver injury in rats and humans upregulates the novel enzyme angiotensin converting enzyme 2. Gut. 2005 Dec;54(12):1790–6. doi: 10.1136/gut.2004.062398. PubMed PMID: 16166274; PubMed Central PMCID: PMCPMC1774784.

22. Zhang R, Wu Y, Zhao M, et al. Role of HIF-1alpha in the regulation ACE and ACE2 expression in hypoxic human pulmonary artery smooth muscle cells. Am J Physiol Lung Cell Mol Physiol. 2009 Oct;297(4):L631–40. doi: 10.1152/ajplung.90415.2008. PubMed PMID: 19592460.

23. Zhang R, Su H, Ma X, et al. MiRNA let-7b promotes the development of hypoxic pulmonary hypertension by targeting ACE2. Am J Physiol Lung Cell Mol Physiol. 2019 Mar 1;316(3):L547–L557. doi: 10.1152/ajplung.00387.2018. PubMed PMID: 30628484.

24. Joshi S, Wollenzien H, Leclerc E, et al. Hypoxic regulation of angiotensin-converting enzyme 2 and Mas receptor in human CD34(+) cells. J Cell Physiol. 2019 Nov;234(11):20420–20431. doi: 10.1002/jcp.28643. PubMed PMID: 30989646; PubMed Central PMCID: PMCPMC6660366.

25. Clarke NE, Belyaev ND, Lambert DW, et al. Epigenetic regulation of angiotensin-converting enzyme 2 (ACE2) by SIRT1 under conditions of cell energy stress. Clin Sci (Lond). 2014 Apr;126(7):507–16. doi: 10.1042/CS20130291. PubMed PMID: 24147777.

26. Asplund A, Ostergren-Lunden G, Camejo G, et al. Hypoxia increases macrophage motility, possibly by decreasing the heparan sulfate proteoglycan biosynthesis. J Leukoc Biol. 2009 Aug;86(2):381–8. doi: 10.1189/jlb.0908536. PubMed PMID: 19401393.

27. Wu F, Wang JY, Chao W, et al. miR-19b targets pulmonary endothelial syndecan-1 following hemorrhagic shock. Sci Rep. 2020 Sep 25;10(1):15811. doi: 10.1038/s41598-020-73021-3. PubMed PMID: 32978505; PubMed Central PMCID: PMCPMC7519668.

28. Taniguchi-Ponciano K, Vadillo E, Mayani H, et al. Increased expression of hypoxia-induced factor 1alpha mRNA and its related genes in myeloid blood cells from critically ill COVID-19 patients. Ann Med. 2021 Dec;53(1):197–207. doi: 10.1080/07853890.2020.1858234. PubMed PMID: 33345622; PubMed Central PMCID: PMCPMC7784832.

29. Crawford KHD, Eguia R, Dingens AS, et al. Protocol and Reagents for Pseudotyping Lentiviral Particles with SARS-CoV-2 Spike Protein for Neutralization Assays. Viruses. 2020 May 6;12(5). doi: 10.3390/v12050513. PubMed PMID: 32384820; PubMed Central PMCID: PMCPMC7291041.

30. Ma D, Chen CB, Jhanji V, et al. Expression of SARS-CoV-2 receptor ACE2 and TMPRSS2 in human primary conjunctival and pterygium cell lines and in mouse cornea. Eye (Lond). 2020 Jul;34(7):1212–1219. doi: 10.1038/s41433-020-0939-4. PubMed PMID: 32382146; PubMed Central PMCID: PMCPMC7205026.

31. Reynolds MR, Singh I, Azad TD, et al. Heparan sulfate proteoglycans mediate Abeta-induced oxidative stress and hypercontractility in cultured vascular smooth muscle cells. Mol Neurodegener. 2016 Jan 22;11:9. doi: 10.1186/s13024-016-0073-8. PubMed PMID: 26801396; PubMed Central PMCID: PMCPMC4722750.

32. Casazza A, Laoui D, Wenes M, et al. Impeding macrophage entry into hypoxic tumor areas by Sema3A/Nrp1 signaling blockade inhibits angiogenesis and restores antitumor immunity. Cancer Cell. 2013 Dec 9;24(6):695–709. doi: 10.1016/j.ccr.2013.11.007. PubMed PMID: 24332039.

33. Lang J, Yang N, Deng J, et al. Inhibition of SARS pseudovirus cell entry by lactoferrin binding to heparan sulfate proteoglycans. PLoS One. 2011;6(8):e23710. doi: 10.1371/journal.pone.0023710. PubMed PMID: 21887302; PubMed Central PMCID: PMCPMC3161750.

34. Appelberg S, Gupta S, Svensson Akusjarvi S, et al. Dysregulation in Akt/mTOR/HIF-1 signaling identified by proteo-transcriptomics of SARS-CoV-2 infected cells. Emerg Microbes Infect. 2020 Dec;9(1):1748–1760. doi: 10.1080/22221751.2020.1799723. PubMed PMID: 32691695; PubMed Central PMCID: PMCPMC7473213.

35. Gan ES, Ooi EE. Oxygen: viral friend or foe? Virol J. 2020 Jul 27;17(1):115. doi: 10.1186/s12985-020-01374-2. PubMed PMID: 32718318; PubMed Central PMCID: PMCPMC7385969.

36. Palazon A, Goldrath AW, Nizet V, et al. HIF transcription factors, inflammation, and immunity. Immunity. 2014 Oct 16;41(4):518–28. doi: 10.1016/j.immuni.2014.09.008. PubMed PMID: 25367569; PubMed Central PMCID: PMCPMC4346319.

37. Zhang Q, Chen CZ, Swaroop M, et al. Heparan sulfate assists SARS-CoV-2 in cell entry and can be targeted by approved drugs in vitro. Cell Discov. 2020 Nov 4;6(1):80. doi: 10.1038/s41421-020-00222-5. PubMed PMID: 33298900; PubMed Central PMCID: PMCPMC7610239.

38. Wing PAC, Keeley TP, Zhuang X, et al. Hypoxic and pharmacological activation of HIF inhibits SARS-CoV-2 infection of lung epithelial cells. Cell Rep. 2021 Apr 5.109020. doi: 10.1016/j.celrep.2021.109020. PubMed PMID: 33852916.

39. Poloznikov AA, Nersisyan SA, Hushpulian DM, et al. HIF Prolyl Hydroxylase Inhibitors for COVID-19 Treatment: Pros and Cons. Front Pharmacol. 2020;11:621054. doi: 10.3389/fphar.2020.621054. PubMed PMID: 33584306; PubMed Central PMCID: PMCPMC7878396.

40. Taylor CT, Colgan SP. Regulation of immunity and inflammation by hypoxia in immunological niches. Nature reviews Immunology. 2017 Dec;17(12):774–785. doi: 10.1038/nri.2017.103. PubMed PMID: 28972206; PubMed Central PMCID: PMCPMC5799081. eng.

41. Rius J, Guma M, Schachtrup C, et al. NF-kappaB links innate immunity to the hypoxic response through transcriptional regulation of HIF-1alpha. Nature. 2008 Jun 5;453(7196):807–11. doi: 10.1038/nature06905. PubMed PMID: 18432192; PubMed Central PMCID: PMCPMC2669289. eng.

42. Merad M, Martin JC. Pathological inflammation in patients with COVID-19: a key role for monocytes and macrophages. Nature reviews Immunology. 2020 Jun;20(6):355–362. doi: 10.1038/s41577-020-0331-4. PubMed PMID: 32376901; PubMed Central PMCID: PMCPMC7201395. eng.

43. Zhu B, Wu Y, Huang S, et al. Uncoupling of macrophage inflammation from self-renewal modulates host recovery from respiratory viral infection. Immunity. 2021 Apr 28. doi: 10.1016/j.immuni.2021.04.001. PubMed PMID: 33951416; eng.

44. Codo AC, Davanzo GG, Monteiro LB, et al. Elevated Glucose Levels Favor SARS-CoV-2 Infection and Monocyte Response through a HIF-1α/Glycolysis-Dependent Axis. Cell metabolism. 2020 Sep 1;32(3):437-446.e5. doi: 10.1016/j.cmet.2020.07.007. PubMed PMID: 32697943; PubMed Central PMCID: PMCPMC7367032. eng.

